# The enduring advantages of the SLOW5 file format for raw nanopore sequencing data

**DOI:** 10.1101/2025.06.30.662478

**Authors:** Hasindu Gamaarachchi, Sasha Jenner, Hiruna Samarakoon, James M. Ferguson, Ira W. Deveson

## Abstract

Nanopore sequencing is a widespread and important method in genomics science. The raw electrical current signal data from a typical nanopore sequencing experiment is large and complex. This can be stored in two alternative file formats that are presently supported: POD5 is a signal data file format used by default on instruments from Oxford Nanopore Technologies (ONT); SLOW5 is an open-source file format originally developed as an alternative to ONT’s previous file format, which was known as FAST5. The choice of format may have important implications for the cost, speed and simplicity of nanopore signal data analysis, management and storage. To inform this choice, we present a comparative evaluation of POD5 vs SLOW5. We conducted benchmarking experiments assessing file size, analysis performance and usability on a variety of different computer architectures. SLOW5 showed superior performance during sequential and non-sequential (random access) file reading on most systems, manifesting in faster, cheaper basecalling and other analysis, and we could find no instance in which POD5 file reading was significantly faster than SLOW5. We demonstrate that SLOW5 file writing is highly parallelisable, thereby meeting the demands of data acquisition on ONT instruments. Our analysis also identified differences in the complexity and stability of the software libraries for SLOW5 (slow5lib) and POD5 (pod5), including a large discrepancy in the number of underlying software dependencies, which may complicate the pod5 compilation process. In summary, many of the advantages originally conceived for SLOW5 remain relevant today, despite the replacement of FAST5 with POD5 as ONT’s core file format.

## INTRODUCTION

Nanopore sequencing is widely used for genome, epigenome and transcriptome analysis[1], and nanopore-based protein sequencing is in development [2]. Its leadership in nanopore development and applications has seen Oxford Nanopore Technologies (ONT) grow into one of the world’s leading genomics companies.

ONT instruments measure the displacement of ionic current as a DNA or RNA molecule passes through a nanoscale protein pore. The resulting time-series current signal data can be translated or ‘basecalled’ into DNA/RNA sequence reads. Raw ONT signal data may also be analysed directly to identify modified DNA or RNA bases [3–5], DNA damage [6], RNA secondary structures [7,8], or for a variety of other purposes [9,10]. Because basecalling models and other analysis algorithms are rapidly evolving, the retention and reanalysis of raw ONT signal data is a common practice.

We previously developed a nanopore signal data file format called SLOW5 (and its binary equivalent BLOW5) [11]. SLOW5 was conceived as an open-source alternative to ONT’s standard data format at the time, called FAST5. SLOW5 addressed limitations in FAST5 and its underlying software library HDF5, which prevented parallel access by multiple CPUs, enabling SLOW5 to achieve order-of-magnitude acceleration of signal data analysis and 20-50% reductions in file size [11]. Such performance enhancements may have profound benefits for users, dramatically reducing the time and costs associated with nanopore data analysis, management and storage. SLOW5 was also a simpler format than FAST5, making it easier both for developers and users to interact with raw ONT signal data [11].

ONT later developed a new signal data file format to replace FAST5, called POD5 (https://github.com/nanoporetech/pod5-file-format). POD5 was designed to fix the design flaws identified in FAST5, enabling faster analysis and smaller file sizes. POD5 is now the signal data file format used by default on ONT sequencing instruments, marking it as a mature format which is suitable for evaluation.

The replacement of FAST5 with POD5 has undoubtedly enhanced ONT as a platform and benefited ONT users. However, SLOW5 was a mature, open-source format available for adoption by ONT prior to the development of POD5. Here we address the question of whether the decision to develop POD5, rather than adopt SLOW5, has resulted in a superior core file format for ONT sequencing. We compare the POD5 and SLOW5 formats, and present benchmarking experiments evaluating file size, analysis performance and usability on a diverse range of computer architectures. Our comparison of POD5 vs SLOW5 is intended to inform nanopore users, developers and commercial vendors on the optimal choice of file format for nanopore signal data.

## RESULTS

### Comparison of POD5 vs SLOW5 file formats

The POD5 and SLOW5 file formats contain an equivalent set of metadata and identical raw signal data. Bi-directional lossless conversion between the two formats is therefore possible. Conversion may be performed using the software package blue-crab (https://github.com/Psy-Fer/blue-crab).

The most obvious difference between the formats is their serialisation or layout. Like most other common genomics file formats (e.g. SAM, FASTA, VCF, BED, etc), SLOW5 is a row-based format. The metadata and all signal data from a given read are stored contiguously and independent reads are laid on a SLOW5 file in sequential rows (**Fig1**; **Table 1**). In contrast, POD5 is column-oriented, with each metadata attribute stored for all reads, followed by signal data for all reads (**Fig1**; **Table 1**). Whereas all signal data points for a given read are adjacent within a SLOW5 file, POD5 stores signal data in sub-reads or ‘chunks’ of a fixed length and chunks from multiple reads may be interspersed (**Fig1**). This layout facilitates writing of multiple reads in parallel, directly to the disk, which is advantageous during data acquisition. However, it means multiple seek operations are required to access all signal chunks and metadata attributes for a given read within the file during data analysis.

**Figure 1.**
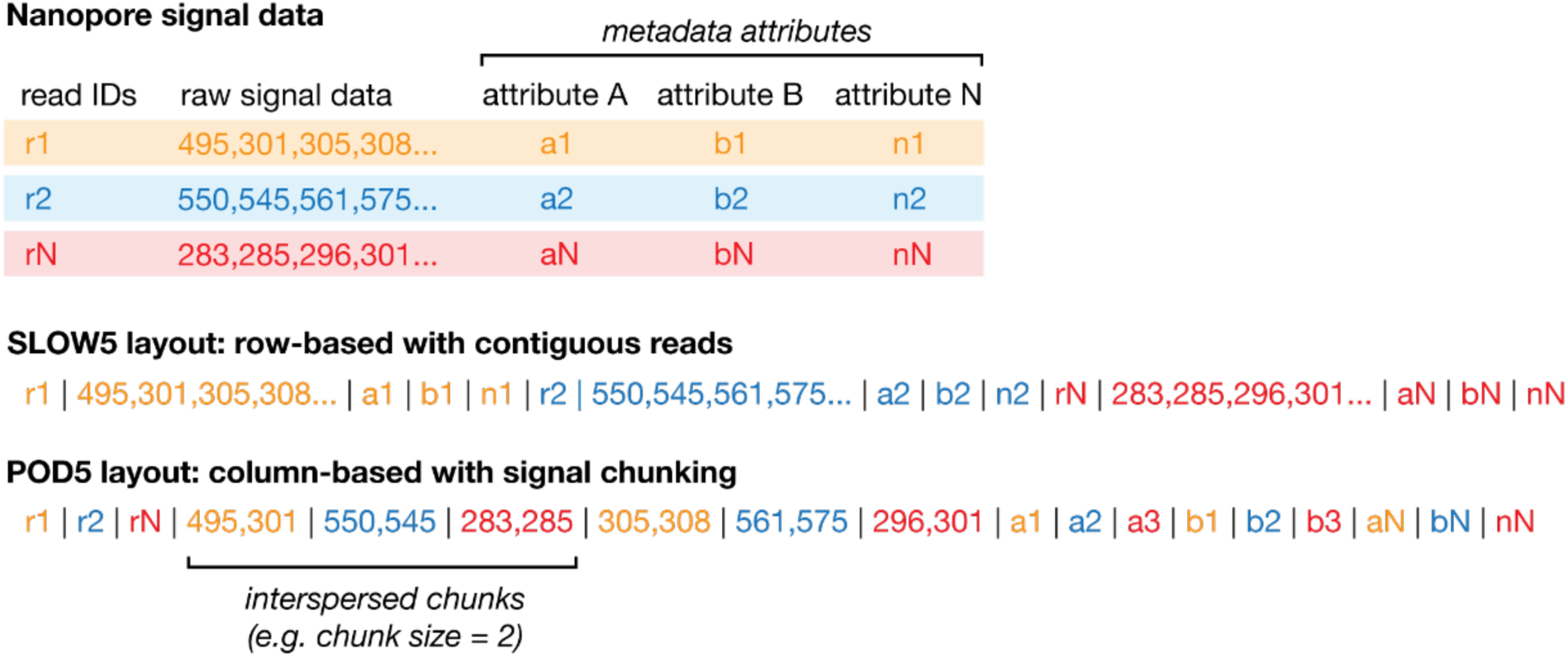
Comparison of SLOW5 vs POD5 file layout. Schematic shows a simplified representation of the layout used to store nanopore signal data in SLOW5 vs POD5 file formats. Upper section shows the basic elements making up a raw nanopore read (three reads shown in different colours): a unique read identifier; a set of metadata attributes describing the read (e.g. channel number, read number, mux values, etc); and the data values for that read. Lower section shows how these elements are serialised onto a linear computer file in SLOW5 format and POD5 format, respectively. SLOW5 uses a row-based layout with all elements for a single read stored contiguously. POD5 uses a column-based layout and additionally stores raw data in ‘chunks’, with chunks from multiple reads allowed to be interspersed within the file. A chunk size of 2 is used here for illustrative purposes, meaning each chunk contains two signal data values (the current default chunk size in POD5 files is 102400).

**Table 1.** Comparison of SLOW5 vs POD5 file formats for nanopore signal data.

| Feature | S/BLOW5 | POD5 |
| --- | --- | --- |
| Layout | Row-based | Column-based |
| Underlying format | None; SLOW5 format written from scratch | Apache Arrow Inter Process Communication (IPC) format; also known as Apache Feather v2 format |
| Variants | ASCII SLOW5 for human readability; binary BLOW5 for computer readability | Binary variant only |
| Signal contiguity | Signal data for each read stored contiguously | Signal data stored in chunks; chunks from a given read may be adjacent or interspersed with chunks from other reads |
| Primary I/O method | Traditional buffered I/O | Memory-mapped (mmap) I/O |
| Accessing a given set of reads | Index file-based | No index file; reads accessed by a 'walker' that traverses through the file |

SLOW5 is a tab-delimited ASCII format with a direct binary equivalent, BLOW5, that does not depend on any other underlying format. POD5 is itself a layer built on top of a primary file format called Apache Arrow Interprocess Communication (IPC) format (**Table 1**). SLOW5/BLOW5 files are read and written by an accompanying library, called slow5lib, which accesses the files through standard C library functions that are very close to the operating system/hardware. POD5 files are read and written by an accompanying library, called pod5, which is built on the Apache Arrow library, which is itself dependent on the library C++17 (**Table 1**). Secondary software libraries for reading/writing SLOW5 and POD5 files are also available in other languages, such as Python and R.

### Benchmarking analysis overview

To evaluate SLOW5 and POD5 formats, we conducted benchmark experiments assessing their basic properties, including file sizes and performance during file reading and writing. We further examined how differences in these properties manifest during real-world analysis scenarios, such as ONT basecalling. We have previously demonstrated that the choice of file format has no impact on analysis outcomes [11,12]. Therefore, we focus here exclusively on performance metrics such as time and memory use, taking equivalent outputs as a given. The binary form of SLOW5 (BLOW5) was used for all experiments. The primary C/C++ libraries for each format were used during all benchmark experiments and secondary libraries in Python, R, Rust or other languages were not considered. Because performance is influenced by the architecture of the computer on which the data is stored and the analysis executed, our benchmark experiments are conducted on an array of different systems, ranging from commercial cloud computing platforms to laptops and a smartphone (**Table 2**). Experiments were run using all available CPU threads on a given system, unless otherwise specified. Finally, we conducted a semi-quantitative assessment of ‘usability’, considering factors such as code complexity, software dependencies and backward/forward compatibility for each file format and its primary software library.

**Table 2.** Overview of computer environments used during benchmarking experiments.

| System name | Description | n x CPU | total cores, cpu threads | RAM | Disk system and file system | O/S |
| --- | --- | --- | --- | --- | --- | --- |
| hpc-ncigadi-cpu | NCI CPU server node | 2 x Intel Xeon Platinum 8274 | 48, 92 | 188 GiB | Lustre distributed file system | Rocky Linux 8.10 |
| hpc-ncigadi-gpu | NCI GPU server node with 8x NVIDIA A100-SXM4-80GB GPUs | 2x AMD EPYC 7742 | 64, 128 | 384 GB | Lustre distributed file system | Rocky Linux 8.10 |
| hpc-pawsey | Pawsey CPU server node | 2 x AMD EPYC 7763 | 128, 256 | 256 GB | Lustre distributed file system | SUSE Linux Enterprise Server 15 SP5 |
| cloud-S3 | Amazon AWS c5a.16xlarge | 1 x AMD EPYC 7R32 | 32, 64 (virtual machine) | 128 GB | s3 bucket (standard tier) mounted using s3fs | Ubuntu 24.04 |
| cloud-EBS | Amazon AWS c5a.16xlarge | 1 x AMD EPYC 7R32 | 32, 64 (virtual machine) | 128 GB | EBS SSD (gp3, 2000GB,IOPS 3000, 125 MB/s) | Ubuntu 24.04 |
| cloud-FsX | Amazon AWS c5a.16xlarge | 1 x AMD EPYC 7R32 | 32, 64 (virtual machine) | 128 GB | fsx for lustre (Persistent SSD, 125 MB/s/TiB, 12TB, no compression, 1500MB/s) | Ubuntu 22.04 |
| server-hdd | Server with HDD | 2 x Intel Xeon Gold 6154 | 36, 72 | 384 GB | 12x10TB HDD drives(RAID6, ext4) | Ubuntu 22.04 |
| server-nfs | Server with an NFS mount | 2 x Intel Xeon Silver 4114 | 20, 40 | 384 GB | NFS mount of a Synology NAS (12 HDD, RAID10, ext4) | Ubuntu 18.04 |
| server-ssd | Server with NVME SSD | 2 x AMD EPYC 7313 | 32, 64 | 512 GiB | 2 x 8TB Mircron 7450 NVME SSD (RAID0, ext4) | Ubuntu 22.04 |
| workstation | A workstation with SSD RAID | 2 x Intel Xeon Platinum 8180 | 56, 112 | 384 GB | 8 x 8TB Micron 5300 SSD(RAID0,ext4) | Ubuntu 20.04 |
| desktop-nvme-ssd | A gaming desktop with NVME SSD | 1 x AMD Ryzen Threadripper 3970X | 32, 64 | 128 GB | 3x 2 TB Samsung SSD 970 EVO Plus NVME SSD drives(RAID0, ext4) | Ubuntu 20.04 |
| desktop-sata-ssd | A desktop with SSD | 1 x AMD Ryzen Threadripper 3970X | 32, 64 | 128 GB | Samsung 870 QVO 8TB SSD (ext4) | Ubuntu 20.04 |
| desktop2-ssd | A desktop with SSD | 1 x 11th Gen Intel Core i7-11700F | 8, 16 | 64 GB | Samsung 870 QVO 4TB SSD (ext4) | Pop!_OS 20.04 |
| desktop3-ssd | A desktop with SSD | 1 x 12th Gen Intel Core i9-12900K | 16, 24 | 64 GB | Kingston SNV2S 2TB NVMe SSD (NTFS) | Ubuntu 22.04 through WSL2 on Windows 11 |
| laptop-nvme-ssd | A laptop with NVME SSD | 1 x 13th Gen Intel Core i9-13900H | 14, 20 | 64 GiB | Samsung 990 Pros 2TB NVMe SSD (NTFS) | Ubuntu 22.04 |
| laptop-usb-ssd | A laptop with an external hard drive | 1 x 13th Gen Intel Core i9-13900H | 14, 20 | 64 GiB | Samsung T7 2TB external USB-c SSD (exFAT) | Ubuntu 22.04 |
| macmini | Apple MacMini external hard drive | 1 x Apple ARM M1 | 8,8 | 8 GB | Samsung T7 2TB external USB-c SSD (exFAT) | macOS 15.1 |
| embedded-system | NVIDIA Jetson Xavier | 1 x ARM v8 | 8, 8 | 16 GB | Samsung 970 EVO Plus 2TB NVMe SSD (ext4) | Ubuntu 20.04 |
| phone | Motorola Moto G54 5G | 1 x MediaTek Dimensity 7020 | 8,8 | 8 GB | Samsung Evo Plus 1 TB microSD (exFAT) | Android 13 |

### File sizes

The series of sequential current signal values making up the body of each read accounts for the majority of the file size in both POD5 and SLOW5 format. Files that store identical signal data are expected to be similar in size. Comparison of identical POD5/BLOW5 files with matched signal compression methods confirmed that any differences in size are negligible (∼0.1% difference; **Table S1**). SLOW5 supports a recently-developed lossless compression strategy *ex-zd* [13], which is not currently supported in POD5. *Ex-zd* compression delivered a modest 2.4% saving in BLOW5 file size compared to the best available POD5 compression (**Table S1**). As reported elsewhere, BLOW5 file sizes may be significantly reduced (∼40%) via *ex-zd* lossy compression with no impact on analysis performance [13], however, this was not considered here, as it is not a like-for-like comparison.

### File reading: sequential access

Sequential access is the simplest and typically the most efficient disk access pattern for reading a file. When reading an ONT signal data file, sequential access means reads are accessed by the disk in the order that they occur in the file (as opposed to random access; see below). This pattern is used during basecalling and DNA/RNA modification calling with ONT’s Dorado software and most other analysis tools where signal reads are handled independently. POD5 uses a technique called ‘mmap’ [14,15] during sequential disk access, whereas SLOW5 uses traditional I/O (**Table 1**).

To evaluate sequential access, we measured the time and memory required to sequentially read a dataset of ∼16 million signal reads in either POD5 or SLOW5 format (hg2_prom_20x; **Table S1**; see **Methods**). The rate of access varied widely depending on the file type and system architecture. However, across all 18 systems, file reading was either equivalent or faster with SLOW5 (**Fig2A**). Equivalent time was required to read a SLOW5 vs POD5 file on some desktop and laptop computers and an Android smartphone, whereas SLOW5 file reading was up to 3.2x faster on commercial cloud, up to 7.8x faster on HPC systems and 2.9x faster on an Apple desktop computer (**Fig2A**). In addition, the memory used during file reading was dramatically higher for POD5 files on all systems, with the exception of the Smartphone (**Fig2B**). The smartphone is an exception here, because it lacks the capacity for mmap and therefore reverts to traditional I/O, which is slower in this context but consumes less memory.

**Figure 2.**
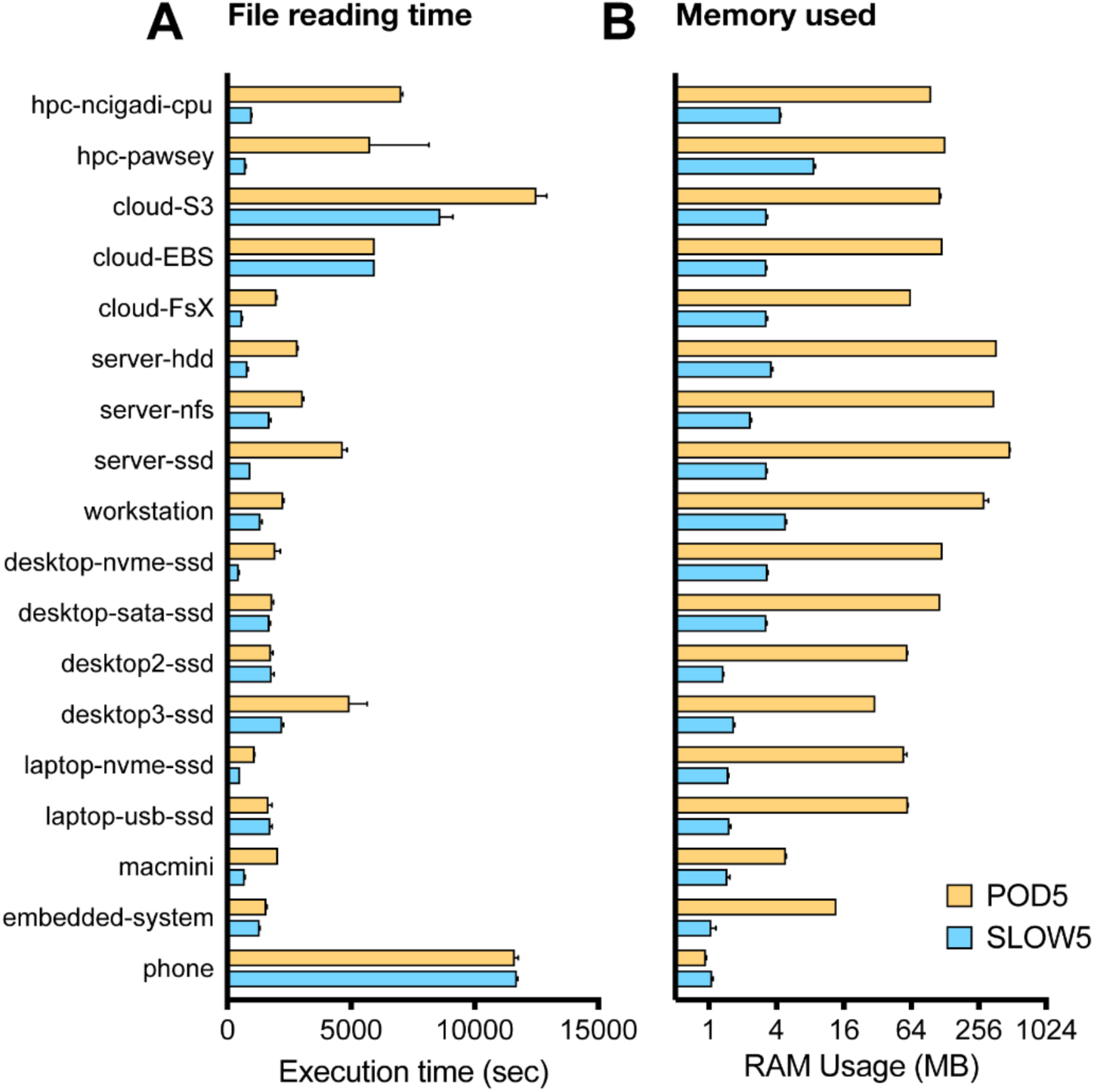
Sequential data access on POD5 vs SLOW5 files. Bar chart shows the time taken (**A**) and memory used (**B**) during file reading in a sequential access pattern for an identical dataset (hg2_prom_20x; see **Table S1**) represented in either POD5 (yellow) or SLOW5 (blue) format. The analysis was performed on 18 different computer architectures (see **Table 2**). Each bar shows the mean of five repeated measurements on each system and error bar shows the range.

Given the large difference in sequential access rates observed on high-performance computing (HPC) and cloud environments, which are commonly used by the genomics community, we next examined how this may influence performance during ONT basecalling. To emulate a typical scenario, we analysed human genome sequencing data from a single ONT PromethION flow cell on an academic HPC system (*hpc-ncigadi-gpu*; **Table 2**; see **Methods**). We ran ONT’s Dorado software on the native POD5 dataset and a SLOW5-enabled version called slow5-dorado on the equivalent SLOW5 file, using the same basecalling model (dna_r10.4.1_e8.2_400bps_hac@v4.2.0), number of threads for file access (8) and file access batch size (1000). We observed a ∼4.7x difference, with SLOW5 basecalling complete within 32 minutes, compared to 2 hours 24 minutes for POD5 (**Table 3**). Memory usage during basecalling was also ∼12.5x higher for the POD5 (374 GB) vs BLOW5 (30 GB) dataset (**Table 3**). Given all other variables were matched, this demonstrates how the more efficient sequential data access of SLOW5 format may manifest in faster basecalling on HPC, cloud and other similar architectures.

**Table 3.**
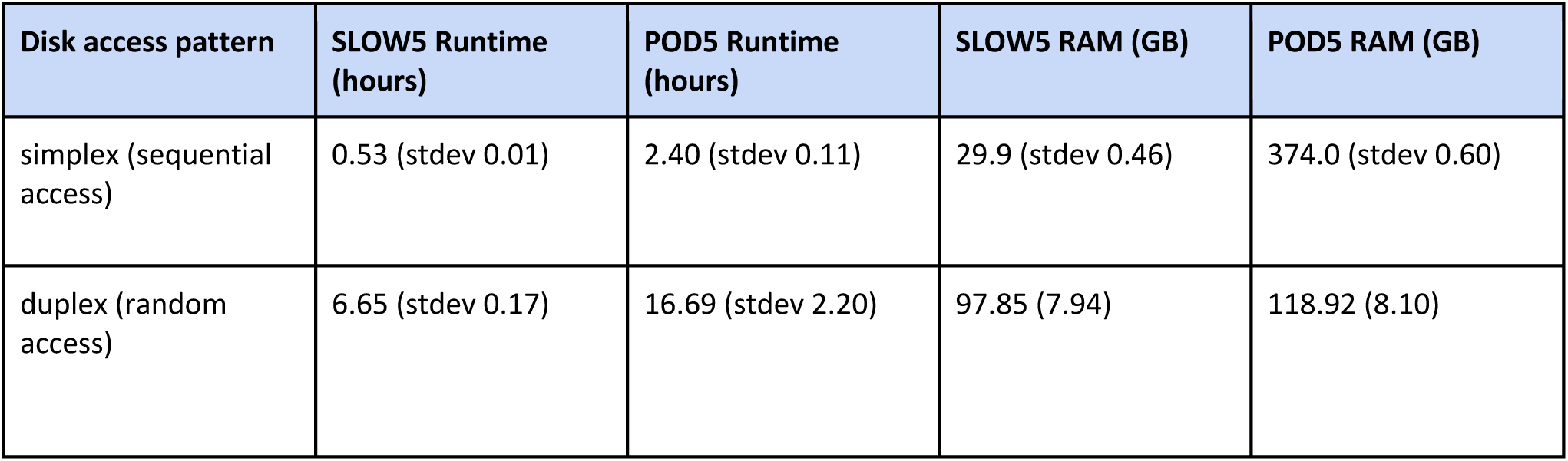
Comparison of simplex and duplex basecalling performance on an HPC system.

| Disk access pattern | SLOW5 Runtime (hours) | POD5 Runtime (hours) | SLOW5 RAM (GB) | POD5 RAM (GB) |
| --- | --- | --- | --- | --- |
| simplex (sequential access) | 0.53 (stdev 0.01) | 2.40 (stdev 0.11) | 29.9 (stdev 0.46) | 374.0 (stdev 0.60) |
| duplex (random access) | 6.65 (stdev 0.17) | 16.69 (stdev 2.20) | 97.85 (7.94) | 118.92 (8.10) |

### File reading: random access

The alternative disk access pattern is random access, where signal reads are accessed non-sequentially from within a file, typically by querying specific reads based on their unique identifiers. While this is more complex than sequential access, it is necessary for any analysis where reads are not handled independently or must be accessed in a specific order. For SLOW5, efficient random access is facilitated by an accompanying index file, which specifies the position of each read within the file [11], whereas POD5 instead uses a random access technique called ‘walker’ (**Table 1**).

We measured the time and memory required to read the same POD5/SLOW5 dataset as above, this time using a random access pattern (see **Methods**). The time taken for file reading again varied between system architectures, however, SLOW5 always showed far superior performance (**Fig3A**). The difference in reading time for SLOW5 vs POD5 ranged from 3.9x on the *cloud-FsX* system to 113x on the *hpc-pawsey* system (**Fig3A**). As with sequential access, the memory used during file reading was higher for POD5 files compared to BLOW5 on all systems, with the exception of the Smartphone (**Fig3B**).

**Figure 3.**
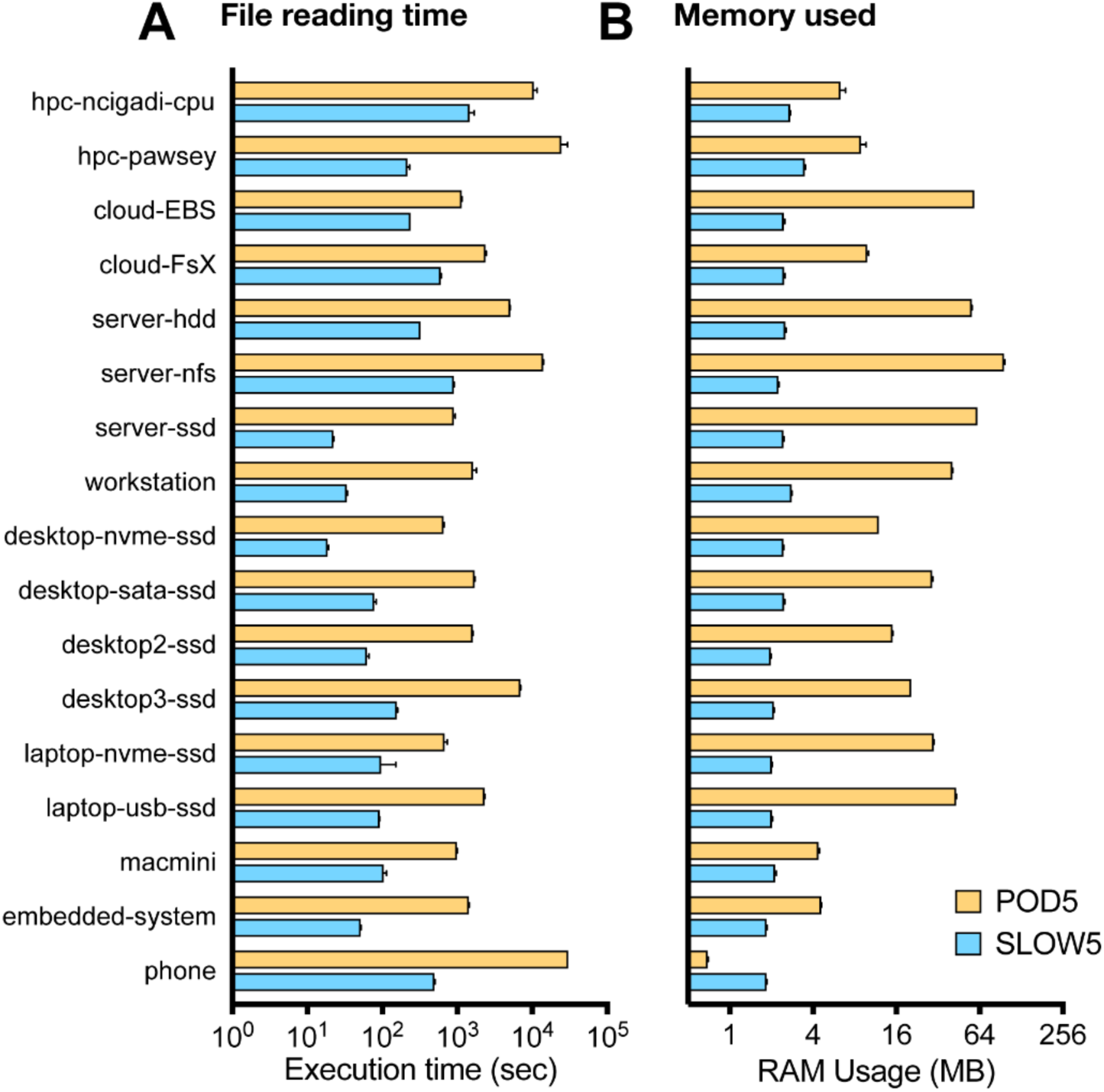
Random data access on POD5 vs SLOW5 files. Bar chart shows the time taken (**A**) and memory used (**B**) during file reading in a random access pattern for an identical dataset (hg2_prom_20x; see **Table S1**) represented in either POD5 (yellow) or SLOW5 (blue) format. The analysis was performed on 17 different computer architectures (see **Table 2**); note the *cloud-S3* system could not be included random access benchmark because the POD5 test were so slow we would be bankrupt by the end of the experiment (not to mention older). Each bar shows the mean of five repeated measurements on each system and error bar shows the range.

Given the large differences observed, we next considered an analysis scenario where random access may impact the overall analysis performance, namely ONT’s duplex basecalling method. During duplex basecalling, independent reads from the complementary strand of a given DNA fragment are analysed together. Because they are often not adjacent within the input file, a random access pattern is preferable for their retrieval. ONT’s Dorado software retrieves complementary signal reads from a POD5 file using its walker strategy, whereas Slow5-dorado uses a file index to find and retrieve complementary reads. We used both tools to run duplex basecalling on matched POD5/SLOW5 files from a single PromethION flow cell, executed on HPC (*hpc-ncigadi-gpu*; see **Methods**). Duplex basecalling with POD5/Dorado took 16.7 hours, compared to 6.7 hours (2.5x improvement) with SLOW5/Slow5-dorado, both running with 8 threads for file access (**Table 3**). Memory usage was again higher for the POD5 (119 GB) vs SLOW5 (98 GB) dataset (**Table 3**). This analysis demonstrates how more efficient file reading for SLOW5 may manifest in large performance improvements for analysis software using random access patterns, such as duplex basecalling.

### SLOW5 writing performance

A potential advantage of the POD5 format lies in its suitability for parallelised data acquisition. An ONT PromethION P48 (ONT’s largest instrument) has 144,000 data channels, meaning it is theoretically possible for 144,000 DNA/RNA molecules to undergo sequencing simultaneously. At even a fraction of this throughput, it is not feasible to store all reads in memory as they are acquired. This is navigated by writing each read to a POD5 output file in sequential chunks, with chunks from multiple reads acquired in parallel being interspersed within the file (**Fig1**).

In contrast, SLOW5 stores all signal data points for a given read contiguously, meaning a new read cannot start being written to the final output file until the previous read is complete. Parallel data acquisition may instead be handled via a two-pass strategy in which reads exceeding a certain size are written in sequential chunks to intermediate files, with one file per read created temporarily. As sequencing is completed for a given read, its intermediate file is read into memory, appended to the combined BLOW5 output file, then the intermediate file is removed. Meanwhile, shorter reads are written directly to BLOW5 output files (see **Methods**).

To evaluate this strategy, we implemented a data simulator, called slowION (https://github.com/hasindu2008/slowION), which mimics data acquisition and reading back (as necessary during live basecalling) from a theoretical nanopore device attached to a given computer (see **Methods**). We used slowION to simulate ONT data acquisition and reading back at increasing scale in order to find the load at which SLOW5 file writing could no longer keep up. We observed the two-pass file writing strategy just described to be highly efficient. Our *workstation* computer, which has comparable specifications to an ONT PromethION, could accommodate up to 189,000 data channels before reaching memory limits and our *desktop-nvme-ssd* computer could accommodate 195,000 channels (**Table 4**). Both computers would therefore be sufficient to accommodate the maximum theoretical data acquisition on a PromethION P48 instrument. Machines with more modest specifications could accommodate fewer channels but were still suitable for relatively high-throughput sequencing. For example, our *laptop-nvme-ssd* computer could accommodate 117,000 parallel channels, sufficient for a PromethION P24 instrument, and our *embedded-system* computer (an NVIDIA Jetson Xavier AGX) could write data for 27,000 channels, theoretically accommodating up to 9x PromethION flow cells on this miniature, low cost chip (**Table 4**).

**Table 4.** Evaluating BLOW5 file writing performance on various computer architectures.

| System name | Max sequencing positions | Max channels | Filesize per hour (GiB) | RAM (GiB) | Reads (Millions) | Samples (Billions) |
| --- | --- | --- | --- | --- | --- | --- |
| server-hdd | 39 | 117,000 | 1434 | 55.7 | 7.7 | 958.0 |
| server-nfs | 25 | 75,000 | 919 | 35.8 | 4.9 | 614.1 |
| server-ssd | 85 | 255,000 | 3125 | 121.0 | 16.8 | 2087.9 |
| workstation | 63 | 189,000 | 23160 | 89.8 | 12.4 | 1547.5 |
| desktop-nvme-ssd | 65 | 195,000 | 2389 | 92.8 | 12.8 | 1596.6 |
| desktop-sata-ssd | 12 | 36,000 | 441 | 17.5 | 2.4 | 294.8 |
| desktop2-ssd | 11 | 33,000 | 404 | 16.1 | 2.2 | 270.2 |
| desktop3-ssd | 10 | 30,000 | 368 | 14.6 | 2.0 | 245.6 |
| laptop-nvme-ssd | 39 | 117,000 | 1433 | 55.7 | 7.7 | 958.0 |
| laptop-usb-ssd | 10 | 30,000 | 368 | 14.6 | 2.0 | 245.6 |
| macmini | 6 | 18,000 | 221 | 2.3 | 1.2 | 147.4 |
| embedded-system | 9 | 27,000 | 331 | 10.9 | 1.8 | 221.1 |

It is not possible for us to directly compare these results to file-writing performance for POD5, because file writing is handled by ONT’s proprietary MinKNOW software. However, this experiment demonstrates that SLOW5 writing performance would not pose a bottleneck at any sequencing scale that is currently feasible.

### Usability

A file format’s ease of use is an important consideration. Although ‘usability’ is context-dependent and inherently subjective, it is possible to assess relevant aspects like the complexity, compatibility and stability of a file format and its accompanying software libraries in a semi-quantitative manner. We attempt to do so here.

The number of dependencies for the primary software libraries for reading/writing SLOW5 (slow5lib) and POD5 (pod5) is an important parameter impacting the ease of installation/compilation and compatibility with different user environments. Slow5lib has only one mandatory external dependency, zlib, which is natively available on any system (**Fig4**). A second dependency, zstd, enables superior signal compression but is optional only. Both zstd and zlib are self-contained, meaning they introduce no secondary dependencies, and can be compiled on any system with common build tools, such as a C compiler and make. In comparison, the pod5 library has nine direct dependencies, including the libraries Apache Arrow, boost and flatbuffers (**Fig4**). These libraries, in particular Apache Arrow, have their own dependencies, giving rise to a large tree of secondary dependencies, which is summarised in **Fig4**. Following this to a branch-depth of eight, we identified 89 unique dependencies for pod5, compared to just three for slow5lib (**Table 5**).

**Figure 4.**
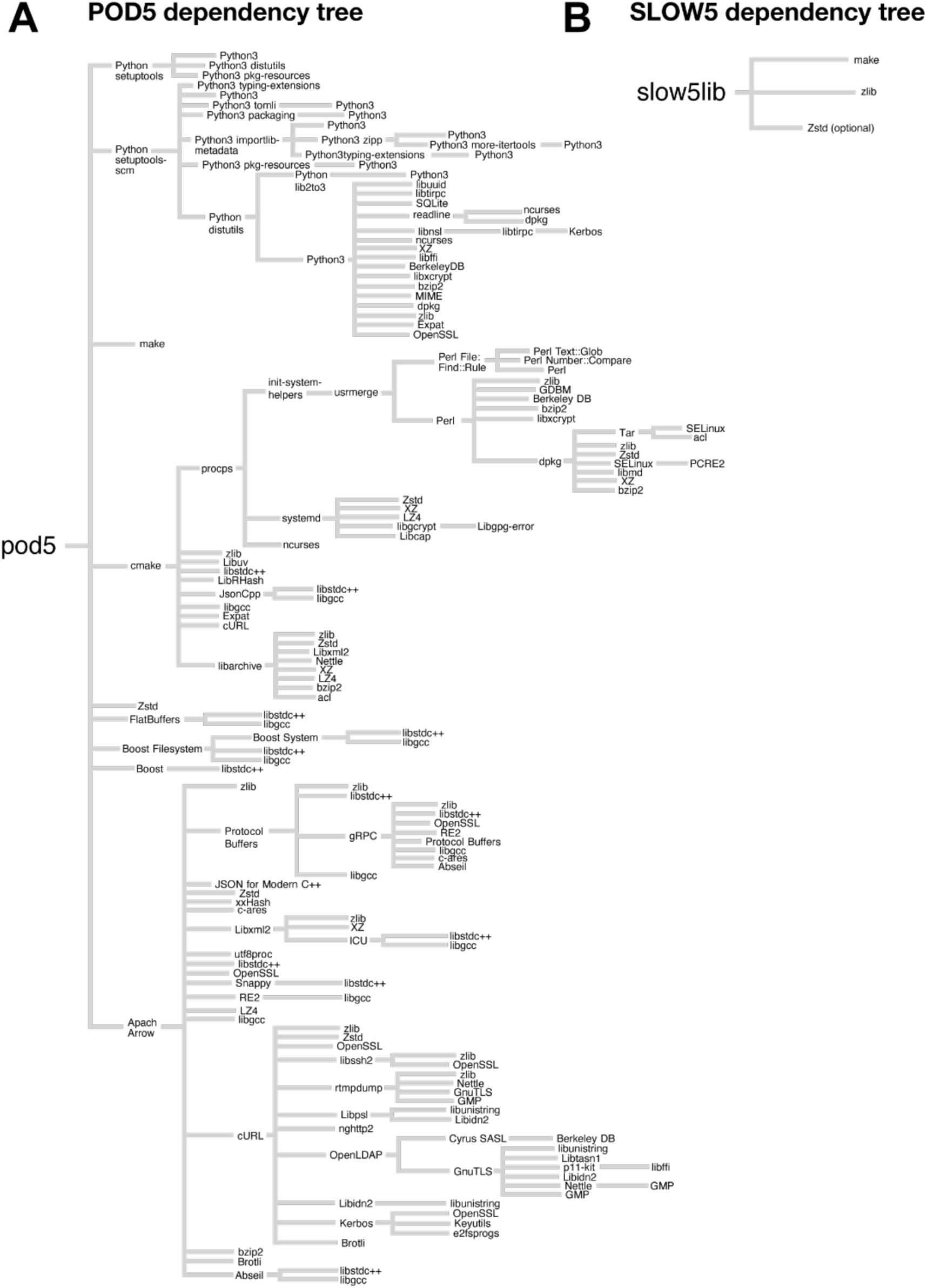
Comparison of POD5 vs SLOW5 software dependencies. The figure depicts the tree of dependencies associated with the POD5 (pod5) and SLOW5 (slow5lib) software libraries. Each dependency subtree is drawn only once; beginning from the trunk, if a subtree is drawn once, it is not drawn again. Some dependency names have been simplified for presentation (e.g., zlib1g-dev is simply zlib and libzstd-dev is just zstd). Standard libc dependency (libc6, not to be confused with libgcc) and the C (e.g., gcc) or C++ (e.g., g++) compilers are not included in the tree. Note that Python dependencies are included for POD5 because these are necessary to generate the version of POD5 for cmake when compiling the POD5 C++ library. The number of unique dependencies identified at each level in the tree is enumerated in **Table 5**.

**Table 5.** Summary of dependency tree for POD5 and SLOW5 libraries. The table enumerates the number of unique software dependencies identified at each level of the dependency trees presented in **Figure 4**. Each dependency was counted only once; i.e. if a given software has already been seen at a higher level in the tree it is not counted again.

| Dependency tree branch-depth | POD5 unique dependencies | BLOW5 unique dependencies |
| --- | --- | --- |
| 1 | 9 | 3 |
| 2 | 40 | 3 |
| 3 | 57 | 3 |
| 4 | 77 | 3 |
| 5 | 82 | 3 |
| 6 | 85 | 3 |
| 7 | 88 | 3 |
| 8 | 89 | 3 |

This array of dependencies complicates the process of compiling the pod5 library, requiring cmake, C++ 17 compiler, conan, ninja-build, etc. To compare the compilation process between slow5lib and pod5, we measured the time taken to download and install all dependencies on a fresh docker image using apt (see **Methods**). For slow5lib this took 26 seconds and produced a static library size of 578 KiB, whereas the POD5 API required 3 minutes and 27 seconds (8.0x longer), with a static library size of 45 MiB (77x larger; **Table 6**). In reality, the compilation process will be influenced by the specifics of the user’s environment. However, this experiment provides a simple measure of the complexity involved, with additional complexity equating to a higher risk of encountering incompatibilities or other barriers.

**Table 6.** Comparison of compilation process for POD5 and SLOW5 libraries. On a fresh docker image of debian:bookworm-slim, the slow5 dependencies were installed from APT and the slow5 library bench branch was compiled with Zstd support. The dependencies for pod5 were downloaded and installed from APT and cmake followed by make was executed.

| Format | Compilation time | shared library size (.so) | static library size (.a) |
| --- | --- | --- | --- |
| SLOW5 | 26 seconds | 0.56 MiB | 1.01 MiB |
| POD5 | 207 seconds | 16.93 MiB | 45.16 MiB |

More specialist software developers, as well as maintainers of the file format itself, may interact directly with the library source code. Here, code complexity is important, with simpler code structure being preferable. We applied a range of standard metrics to evaluate the code bases for slow5lib and the pod5 library, excluding their dependencies (**Table 7**; see **Methods**). The pod5 code is larger (4.1x more lines), has higher cyclomatic complexity (CCM; 1.27x), contains a larger number of parameters (1.78x) and functions (4.3x). The only metric on which slow5lib was more complex was its higher CCM per function (1.3x).

**Table 7:** Source code complexity metrics for the SLOW5 and POD5 code base. LOC (lines of code without comments), CCN (cyclomatic complexity number summed over all functions), Tokens (number of tokens - a token is the smallest unit in a program, e.g., operators, keywords, identifiers, separators), Params (total function parameters), Fns (number of functions) and per function averaged statistics. Only includes the main GitHub repository without any dependencies.

| Format | LOC | CCN | Tokens | Params | Fns | Avg.LOC/Fn | Avg.CCN/Fn | Avg.Tokens/Fn |
| --- | --- | --- | --- | --- | --- | --- | --- | --- |
| slow5 | 6916 | 1372 | 39184 | 649 | 252 | 10.2 | 2.1 | 82.0 |
| pod5 | 28144 | 1749 | 59362 | 1153 | 1072 | 10.7 | 1.6 | 79.1 |

Finally, we assessed the stability or backward/forward compatibility of each code base, which is another important variable for developers. To do so we generated a POD5 and SLOW5 file using every historic version release of the pod5 library (46 versions) or slow5lib (14 versions; **Fig5A**) then attempted to read each file with every other library version, creating a file/library compatibility matrix displayed in **Fig5B** (see **Methods**). For SLOW5, there were no breaks in backward compatibility and a single change in the file format at version 0.3.0, where forward compatibility was broken (**Fig5B**). For POD5, there was a series of breaks in both backward and forward compatibility, meaning that some previous file versions cannot be read with current library versions, and vice versa (**Fig5B**). Next, we assessed the code base of each library to identify breaking changes in successive version releases, i.e. where compatibility is disrupted by a change in the library or command line tool, requiring intervention from the user to restore compatibility (see **Methods**). While there were no such breaking changes in the history of slow5lib, we identified 5 for pod5, although notably no breaking changes were made since April 2023 (version 0.1.12), indicating good stability from this point onwards (**Fig5A**).

**Figure 5.**
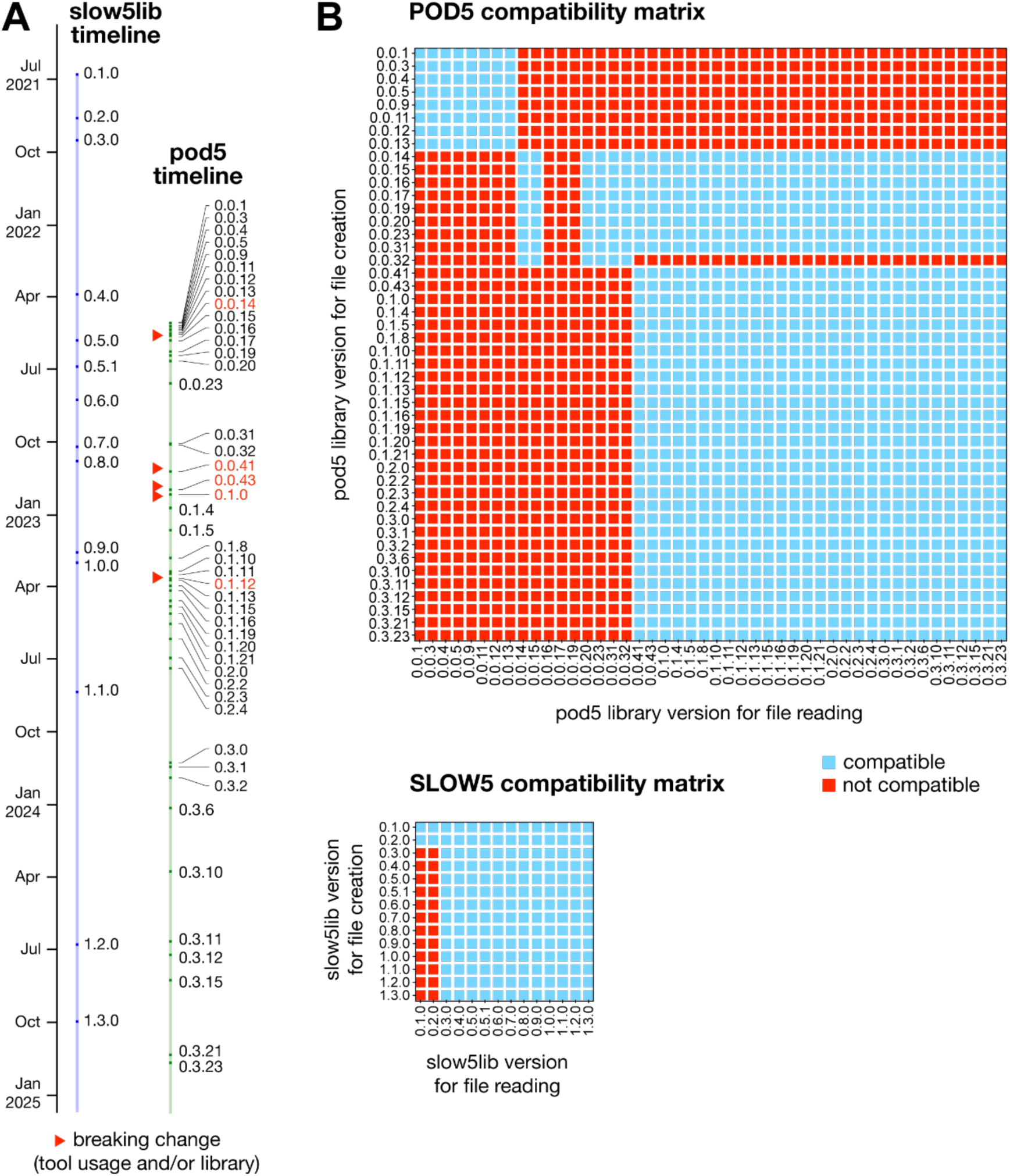
Historical stability of POD5 and SLOW5 formats. (**A**) The figure depicts a timeline of version releases for the POD5 (pod5) and SLOW5 (slow5lib) software libraries. Red markers indicate releases which introduced a ‘breaking change’, i.e. where compatibility is disrupted by a change in the library or command line tool, requiring intervention from the user to restore compatibility. (**B**) The upper figure shows the compatibility of a POD5 file generated with every previous pod5 library version to be read with every other version. Blue squares indicate the file could be read (compatible) and red squares indicate the file could not be read (not compatible). The lower figure presents the equivalent matrix for reading SLOW5 files with all previous slow5lib version releases.

## DISCUSSION

Here we attempt to determine the optimal format for storing raw signal data from nanopore sequencing experiments. Our comparative evaluation of ONT’s POD5 file format and the open-source alternative, SLOW5, identified several areas where SLOW5 showed clear advantages. File reading was faster for SLOW5 on most computer architectures when using a sequential access pattern, faster on all systems when using a random access pattern, and SLOW5 reading used considerably less memory in all scenarios. This manifested in faster, cheaper ONT basecalling with both simplex (sequential access) and duplex (random access) methods. Despite testing on a wide variety of different systems, we could find no instance in which POD5 file reading was significantly faster than SLOW5.

The advantages of SLOW5 for sequential access are the result of its simple row-based layout with contiguous reads, as opposed to the column-oriented and chunk-based layout used in POD5, which necessitates multiple seek operations to retrieve a single read. While the time penalty for performing these seek operations is smaller on systems with fast SSD storage, there is never a scenario where this process could be faster than simply reading sequential rows via traditional I/O, as for SLOW5. Reading a file in a random access pattern is known to be slower than sequential access. However, this is necessary for any analysis where reads are retrieved in a specific order (e.g. based on their genome coordinates). SLOW5 uses a file index to facilitate efficient random access. This is sufficiently fast to enable semi-interactive browsing of signal data aligned to a given genome or transcriptome region using *Squigualiser* [16] and can even be used to extract specified reads from large files on a remote server using *slow5curl* [17]. A comparable indexing strategy is not supported for POD5. This makes it particularly poorly suited for random access, ranging from 3.9x to 113x slower than SLOW5 on the architectures tested here.

While the complex layout used in POD5 creates challenges during file reading, it has supposed advantages for file writing. By storing reads in chunks that may be either adjacent or interspersed in the file, multiple reads can be directly written to the file effectively simultaneously. This parallelisation of data acquisition is necessary to support high-volume sequencing on an ONT instrument. However, we show here that this requirement can also be fulfilled via a two-pass file writing strategy, which enables highly parallel data acquisition with SLOW5. Our slowION simulation indicates that even a laptop computer can perform SLOW5 file writing at sufficient capacity to accommodate an ONT PromethION P24 instrument at maximum sequencing throughput. SlowION is the simplest implementation of this file-writing strategy, shown here as a proof-of-concept. With further optimisation, such as multi-threading, we anticipate the limits of file writing achievable with SLOW5 are likely much higher. An added benefit of the two-pass approach is that reads are naturally separated by channels and laid onto the final output file in order they were sequenced. This is beneficial when identifying and processing complementary reads during duplex basecalling, which are typically sequenced sequentially on a given channel. Overall, although the chunk-based strategy used in POD5 is a valid solution to the challenge of parallel data acquisition, SLOW5 is similarly suitable to handle the demands of file writing on ONT instruments.

Our semi-quantitative usability analysis identifies some important differences between POD5, SLOW5 and their accompanying libraries. Most notably, the pod5 library has a large tree of direct and secondary dependencies, largely arising from its use of the Apache Arrow software library. In contrast, slow5lib has only three direct dependencies – one of which is optional and the other two ubiquitous – and no secondary dependencies. This ensures slow5lib can be easily compiled on any Linux system, whereas pod5 compilation is more likely to encounter conflicts. To alleviate this, ONT provides pre-compiled pod5 binaries for many common architectures. This is sufficient for most users but cannot cover all possible architectures. Conversely, pod5 binaries are available for Windows systems, whereas slow5lib is not currently supported on Windows. This does not reflect an inherent limitation of the SLOW5 format. Rather, Windows development has not been pursued because Windows Subsystem for Linux (WSL) and other similar solutions are popular among the bioinformatics community, while direct Windows users are relatively rare (if you are Windows user who wants to use SLOW5, please contact us).

In summary, we found SLOW5 to have multiple advantages in terms of performance and usability, and few – if any – parameters on which POD5 was preferable. While ONT instruments do not output data in SLOW5 format directly, users can perform lossless POD5-to-SLOW5 conversion using the software package blue-crab (https://github.com/Psy-Fer/blue-crab). We continue to maintain SLOW5 and a range of associated software as a community-centric alternative for ONT signal data storage and analysis: https://hasindu2008.github.io/slow5/

## METHODS

### File reading benchmarks

#### Matching conditions

All benchmark experiments were performed on the binary representation of SLOW5, known as BLOW5. This is similar to POD5, which has only a binary version. Performance benchmarks were performed on a large number of different computer systems, over an extended period of time. The pod5 library version 0.3.2 was selected for use in all benchmark experiments and slow5lib version 1.2.0 was used after patching to match all conditions with the pod5 library (see below). These were the latest library versions available at the time of commencement. ONT developers confirmed that the POD5 file format was stable at that point in time and therefore no major performance changes were anticipated in the future.

For a fair comparison, it is essential to match all possible conditions, including the compiler versions, compiler optimisation flags and the compression methods. We used pre-compiled binaries for pod5, provided by ONT, and deduced the compiler versions, compiler optimisation flags, and the zstd library version used in those compiled binaries to be gcc/g++ 10, -O3 optimisation level and zstd 1.5, respectively. We therefore compiled slow5lib with those settings to ensure matched conditions. POD5 implemented a newer version of VBZ compression that uses an SIMD-accelerated 2-byte variant of StreamVByte, as opposed to the 4-byte variant of StreamVByte used in slow5lib. We therefore ported this newer VBZ into slow5lib for the purposes of this benchmark. Details of all the commands/conditions that are matched are outlined in **Supplementary Note 2**.

#### Benchmark programmes

The benchmark programmes are written in the C/C++ programming language. These benchmark programmes perform the following steps repeatedly for a batch of reads at a time.

1) Read the data from the disk and perform decompression, and parse the binary data into memory structures.
2) Perform a simple summation of the signal data values.
3) Output this sum along with metadata.

The time for Step 1 was measured by using the “gettimeofday” function called just before and after Step 1. Steps 2 and 3 mimic a real programme, such as basecalling that consumes loaded data, mainly to prevent compilers from optimising any unused portions in the code. The peak RAM usage during the lifetime of the benchmark programme was measured by using the *maxrss* value from the “getrusage” function. The source of the benchmark programme code can be found in the slow5-pod5-bench repository: https://github.com/hasindu2008/slow5-pod5-bench.

The code for reading POD5 for sequential access was directly adapted from code used in ONT Dorado for simplex basecalling (DataLoader::load_pod5_reads_from_file in Dorado v0.7). This includes the thread model used in Dorado for accessing POD5 files with multiple threads. The data fields accessed and the order of access to the fields are the same as in Dorado (see **Supplementary Note 1**). To ensure a fair comparison, we used the same thread model to access the same data fields in the same order in BLOW5, even though this order of access is not the order in which the data is laid out in a BLOW5 file (and therefore not the most efficient way to read a BLOW5 file). Furthermore, when populating data memory structures, POD5 datatypes are unchanged whereas BLOW5 datatypes are converted to match those in POD5, for which the conversion penalty is also included in time measurements for BLOW5. Some examples are:

- sampling_rate which is a double in BLOW5 is converted to uint16_t to match POD5
- channel_number which is a string in BLOW5 is converted to uint16_t to match POD5
- The fields range and digitisation in BLOW5 are both read and are divided to compute the single “scale” value in POD5.

The ‘walker’ strategy for performing random access to reads was also directly adapted from code used in ONT Dorado for duplex basecalling (DataLoader::load_pod5_reads_from_file_by_read_ids function in Dorado v0.7). The sanity of our code implementation for POD5 was verified by POD5 developers (for example, see https://github.com/nanoporetech/pod5-file-format/issues/136).

#### Benchmarking procedure

The BLOW5 files were first converted to the VBZ compression to match that in POD5 (see above). For random access benchmarks, a list of 500,000 readIDs was randomly created, with the same list being used for both POD5 and BLOW5. Benchmark programmes are compiled with g++10 with -O3 optimisation (see above).

We performed the benchmarks on various systems (**Table 2**) and datasets (**Table S1**). Benchmark experiments were executed using all hardware CPU threads available on the system. A read batch size of 1000 was used in all cases, as this value is hardcoded in POD5 and thus can’t be changed. Each experiment was executed 5 times, and the average execution time and RAM usage were recorded, as described above.

Mitigating the confoudning impact of the operating system disk cache is crucial for a file format benchmark. To do so, we used datasets that are larger than the amount of RAM available on the systems (**Table S1**) and, additionally, clean the operating system disk caches (pagecache, dentries and inodes) before every test. In Linux systems where we have root access, this was done by writing the file “/proc/sys/vm/drop_caches”. On Mac we used the “purge” command. On systems without root access, such as academic supercomputers and the Android smartphone, we write and then read large amounts of data (larger than the system RAM available) to fill the disk caches with this mock data.

Example commands are provided in **Supplementary Note 2**, and benchmarking scripts are available in the slow5-pod5-bench repository: https://github.com/hasindu2008/slow5-pod5-bench.

#### Basecaller benchmarks

For basecalling, slow5-dorado v0.3.4 was used for BLOW5 and dorado 0.3.4 for POD5. For simplex basecalling, the basecalling was performed with the high-accuracy model on the dataset hg2_prom_20x. For duplex basecalling, the super-accuracy model was used on hg2_prom_duplex dataset. All basecalling experiments were performed on the hpc-ncigadi-gpu academic supercomputer with 8 NVIDIA A100 GPUs. As we used prebuilt binaries, the conditions mentioned above are not perfectly matched. However, the conditions favour POD5, as we used the less efficient streamVByte variant, older compiler versions and less aggressive optimisation flags in slow5-dorado binaries. Furthermore, in ONT Dorado (both simplex and duplex basecalling), the number of threads used for POD5 reading is hardcoded to the number of GPUs available on the system (8). Thus, we have forced slow5-dorado also to use 8 threads for BLOW5 access. The BLOW5 batch size is also made 1000 to match the hardcoded value in POD5. As for the previous benchmarks, we cleaned the disk caches before each experiment, and the average across 5 executions is taken. Commands are provided in **Supplementary Note 2**.

### SLOW5 writing benchmark

We implemented slowION to simulate the data acquisition process using SLOW5 files, including writing data to disk and subsequently reading it back to mimic live basecalling. slowION is implemented using C on top of the slow5lib library. slowION spawns three distinct thread pools, each with a number of threads equal to the number of sequencing positions, implemented using POSIX threads:

1. **Acquisition threads** simulate data acquisition. Read signals shorter than a predefined length threshold (the chunk size) are written directly to a BLOW5 file. Each thread writes to a separate file, so one BLOW5 file is created per position. Signal chunks of longer reads (exceeding the chunk size) are written to temporary binary files, with one file created per read.
2. **Merge threads** monitor for completed reads in temporary binary files, read them, append them to a separate BLOW5 file and delete the temporary files. Here, one BLOW5 is created for each thread (sequencing position) and is separate to the files used by threads in Step 1 above.
3. **Read-back threads** simulate live basecalling by reading completed reads from BLOW5 files. Again one thread is used per sequencing position, which is responsible for the two respective BLOW5 files generated from Steps 1 and 2 above.

If any thread in any pool fails to complete its task within the real-time constraint, a warning is generated. To ensure that data is truly written to disk (and not just cached in memory), the simulation is run for 1 hour on each system, with total data volume far exceeding available RAM. We repeated the simulation while incrementally increasing the number of sequencing positions in slowION until warnings begin to appear. By gradually increasing the number of simulated positions in parallel, we identify the point at which the first warning appears (at least one warning). This is considered the ceiling for parallel data acquisition on a given system. The occurrence of even a single warning is treated as a failure under this strictest evaluation criteria, although in practice, a system can tolerate such incidental warnings. The exact commands used are provided in **Supplementary Note 2**.

To understand the process, consider the following example:

Using a chunk size of 200,000 signal values, which are 2 bytes each, a PromethION with 144,000 data channels translates to ∼54 GiB of memory. This is comfortably accommodated in the RAM of a PromethION compute tower. Given a 5kHz sampling rate and 400bps translocation speed, 200,000 signal samples translates to ∼16 Kbases. In a given sequencing run, a lot of reads are shorter than 16Kbases, and therefore fit inside a single chunk. Such reads are first cached in the RAM, then directly written to a BLOW5 file. Only the reads longer than this threshold will go through two-pass file writing. These longer reads are first written to an intermediate temporary binary file, with one temporary file created for each read. As soon as the acquisition of a long read is completed, the temporary file is read into memory, appended to a BLOW5 file and the temporary file is removed.

### Evaluating usability

#### Dependencies and compilation

To create the software dependency trees for the slow5lib C library and the pod5 C/C++ library, the direct dependencies were first identified from the source code and documentation. For each direct dependency identified, the *apt info* command on Debian 12 (Bookworm) was then executed, and the “Depends” tag of the output was used to identify its dependencies, forming the second layer in the dependency tree. This was repeated for all the dependency levels until we reached a package with no further dependencies. When counting the number of dependencies, we only counted unique software – i.e. for a given level, we do not recount a dependency if it has been seen at a previous level of the tree.

The time for building the slow5 C library (with ztsd support) and pod5 C/C++ libraries was separately measured by performing the building steps inside fresh Docker instances. The experiments were performed on the server-nfs system (**Table 1**) using Debian 12 (Bookworm) as the Docker image. The execution time for the steps (including the installation of dependencies using the apt package manager) was measured using the GNU time utility. Sizes of the generated library files were measured using the *du* command. Further details are provided in **Supplementary Note 2**.

#### Source code complexity

Source code complexity metrics were measured using lizard (https://github.com/terryyin/lizard) version 1.17.17. For source code complexity measurements, only the source code files inside the GitHub repository required to build the C/C++ library were included. External dependencies, documentation, tests, benchmarks and examples, and Python wrappers were not included. Comments and white space were not included for line counts. Further details are provided in **Supplementary Note 2**.

#### Compatibility and stability

To evaluate the stability and backwards/forwards compatibility of POD5 and SLOW5, we developed a simple ‘test program’ that sums up the picoampere converted raw signal for SLOW5 and POD5, separately. We then prepared a BASH script that iteratively tests the compatibility of a given library version with a file created by a different version. The testing process is as follows:

1. Install all software library versions. For SLOW5, we downloaded the slow5lib source from GitHub and compiled it. For POD5, we used pre-compiled library binaries downloaded from GitHub .
2. Install all toolkit versions. For slow5tools, we downloaded binaries from the slow5tools GitHub respository. For pod5, we installed the pod5 toolkit python package.
3. Create a POD5/BLOW5 file from each installed version of the relevant toolkit
4. Compile the ‘test program’ against each library version
5. Run each ‘test program’ created in Step 4 above with all the files created in Step 3 above, then check whether the output of the test program is correct.

If the output of the program was correct for a given combination of versions, those versions were deemed to be compatible. If the program failed to run or produced an output that was incorrect, those versions were deemed to be incompatible. Some versions of pod5 performs an internal conversion of one version to another on the disk as a temporary file before reading it; we do not consider this as a failure because it is handled internally. In the version release timeline, we consider a breaking change to have occurred when the script or the test programme code had to be modified to work with a given version due to a change in the toolkit command or the C library API. We did not include any cases where the binaries/source code for a particular version was missing in the pod5 repository.

### Blue-crab implementation

A python tool, blue-crab, was developed to allow POD5-to-S/BLOW5 and S/BLOW5-to-POD5 lossless conversion. Using the pod5 and pyslow5 python libraries, blue-crab can convert many split files of one format to a mirrored folder structure of the other format, or many files to one file. It can be installed with a ‘pip install blue-crab’ command. Extensive testing is carried out to ensure compatibility is maintained with new releases as the POD5 schema changes over time.

## ACKNOWLEDGEMENTS

We acknowledge the following funding support from Australian Medical Research Futures Fund grants MRF1173594, and MRF2023126 (to I.W.D.), Australian Research Council DECRA Fellowship DE230100178 and Australian Research Council’s Discovery Project DP230100651 (to H.G). Resources from the National Computational Infrastructure (NCI Australia) and Pawsey Supercomputing Research Centre were used for this work. The views expressed herein are those of the authors and are not necessarily those of the funding bodies.

## AUTHOR CONTRIBUTIONS

H.G. and I.W.D. devised the experiments and prepared the manuscript, with support from all authors. H.G. and H.S. implemented the benchmark programmes. J.F. designed and implemented blue-crab. S.J. implemented the new VBZ compression used in POD5 to BLOW5. H.G., S.J. and H.S. performed the benchmark experiments. All authors read and approved the final manuscript.

## DECLARATIONS

I.W.D. manages a fee-for-service sequencing facility at the Garvan Institute of Medical Research and is a customer of Oxford Nanopore Technologies but has no further financial relationship. H.G., J.M.F. and I.W.D. have previously received travel and accommodation expenses from Oxford Nanopore Technologies. H.G. has a paid consultant role with Swan Genomics PTY. The authors declare no other competing financial or nonfinancial interests.

## DATA & CODE AVAILABILITY

Benchmark code: https://github.com/hasindu2008/slow5-pod5-bench Bluecrab: https://github.com/Psy-Fer/blue-crab

slowION: https://github.com/hasindu2008/slowion

The BLOW5 files (zlib+svb-zd compression) of the data are available through the ENA: hg2_prom_20x [ERR12997167], hg2_prom_40x [ERR12997168], hg2_prom_duplex[ERR13475640], uhr_rna_prom [ERR12997170]. These BLOW5 files can be converted to BLOW5 (vbz) and POD5 versions as explained at the **Supplementary Note 2**.

## SUPPLEMENTARY INFORMATION

**Table S1.** Summary of datasets used during benchmarking experiments.

| Dataset name | Description | N reads | Total signal Samples | BLOW5 (VBZ) Size | BLOW index size | POD5 (VBZ) Size | BLOW5 (ex-zd) size |
| --- | --- | --- | --- | --- | --- | --- | --- |
| hg2_prom_20x | HG002 PromethIO N LSK114 at 5kHz | 16.1 million | 903.2 billion | 739 GiB | 0.8 GiB | 740 GiB | 723 GiB |
| hg2_prom_40x | HG002 PromethIO N LSK114 at 5kHz | 21.7 million | 2035.7 billion | 1659 GiB | 1.1 Gib | 1661 GiB | 1622 GiB |
| hg2_prom_duplex | HG002 PromethIO N LSK114 'high duplex' at 5kHz | 4.9 million | 895.4 billion | 731 GiB | 0.2 GiB | 733 GiB | 716 GiB |
| uhr_rna_prom | Universal human RNA; direct RNA004 | 16.4 million | 641.1 billion | 549 GiB | 0.8 GiB | 549 GiB | 539 GiB |

### Supplementary Note 1: Field access order

The fields accessed (in order) are as below:

1. run_acquisition_start_time_ms
2. sample_rate
3. read_id
4. num_samples
5. raw_signal
6. start_sample
7. calibration_scale (scaling)
8. calibration_offset (offset)
9. read_number
10. well (mux)
11. channel (channel number)
12. acquisition_id (run_id)
13. flowcell_id
14. sequencer_position (position_id)
15. experiment_name (experiment_id)

This is the order used in Dorado v0.7 (https://github.com/nanoporetech/dorado/blob/9ac85c65fc873a956bda00b2f5608b2bf72d9e7c/dorado/da ta_loader/DataLoader.cpp#L876)

### Supplementary Note 2: Detailed Methods

#### Format/compression conversion

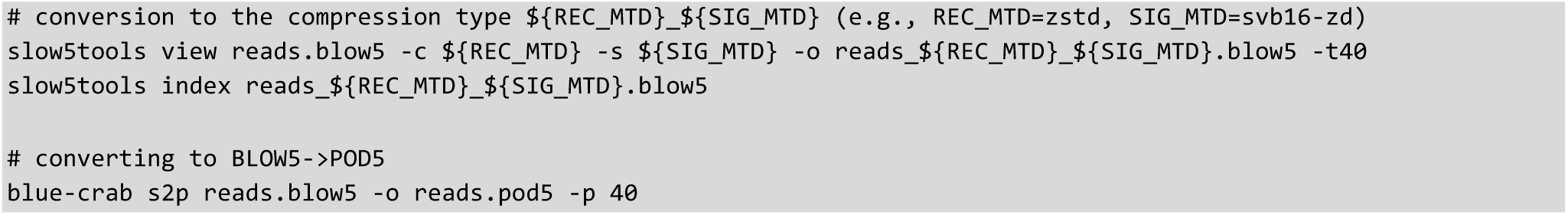

Versions:

- for zstd+svb16-zd (vbz): slow5tools vbz branch [https://github.com/hasindu2008/slow5tools/tree/vbz commit 8a366bf6dffe0c94fd0ec148cca22f09e47c31e5].
- zstd+svb-zd and zstd+ex-zd: slow5tools 1.3.0
- blue-crab 0.1.0

#### File size measurements

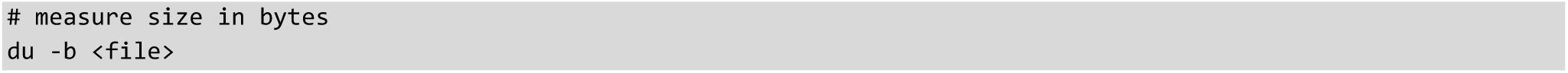

#### Disk cache cleaning

- For systems with root access: https://github.com/hasindu2008/biorand/blob/master/clean_fscache.c
- For systems without root access: https://github.com/hasindu2008/biorand/blob/master/clean_fscache2.c

#### Reading

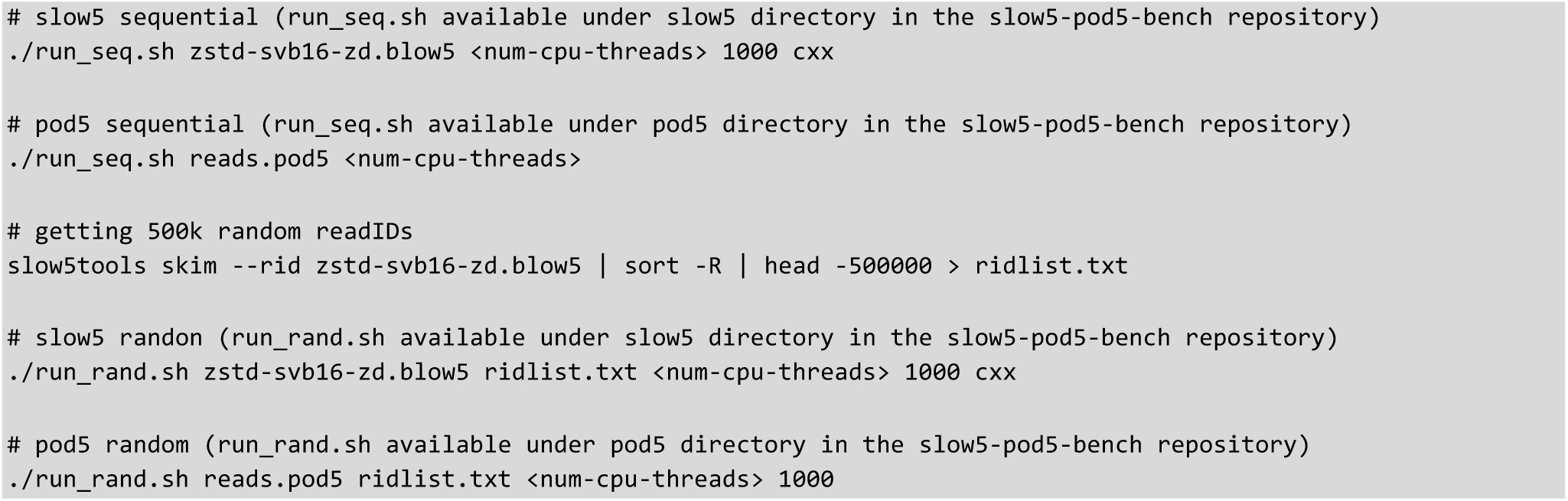

#### Basecalling

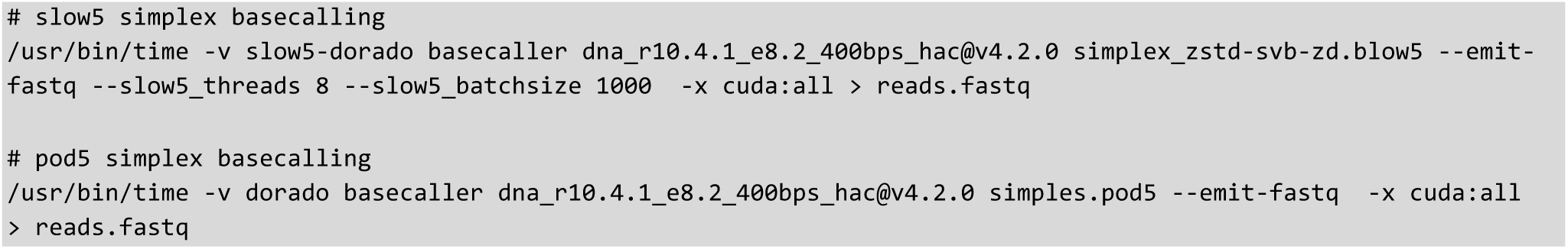

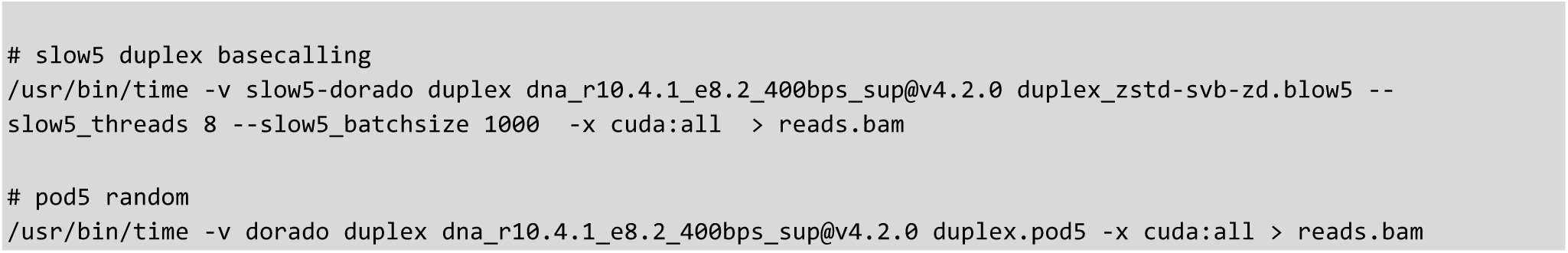

Versions:

- slow5-dorado v0.3.4
- Dorado v0.3.4

#### Writing

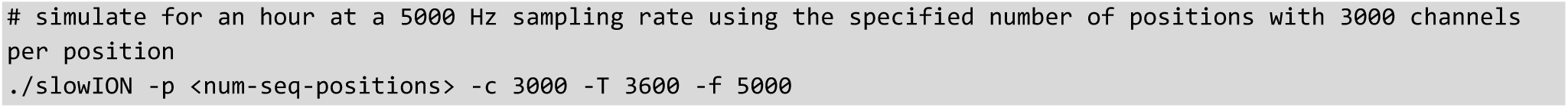

#### Versions

- slowION v0.1.0

#### Dependency tree

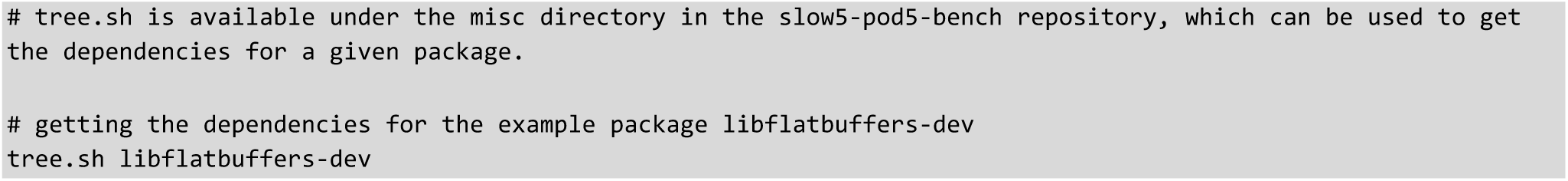

versions:

- slow5lib v1.2.0
- pod5 v0.3.2
- Docker version 26.1.3; *debian:bookworm-slim* image (Debian 12)

#### Notes

- When drawing the dependency tree, dependency names have been simplified (e.g., zlib1g-dev is simply zlib and libzstd-dev is just zstd).
- Standard libc dependency (*libc6*, not to be confused with libgcc) is not included in the tree. This is because *libc6* is at the leaf of almost every library. For instance, though zlib and zstd have *libc6* at the end, it is not drawn.
- Each dependency subtree is drawn only once. We begin from the bottom, and if a subtree is drawn once, it is not drawn again. As an example, *libstdc++* under Abseil has *libgcc* dependency listed, but above this, any other libstdc++ nodes do not have this libgcc.
- The C (e.g., gcc) or C++ (e.g., g++) compilers are not included in any of the trees.
- Note that Python dependencies are included for POD5, not because we included the Python wrapper for POD5, but because these are necessary to generate the version of POD5 for cmake when compiling the POD5 C++ library.

#### Source code compilation

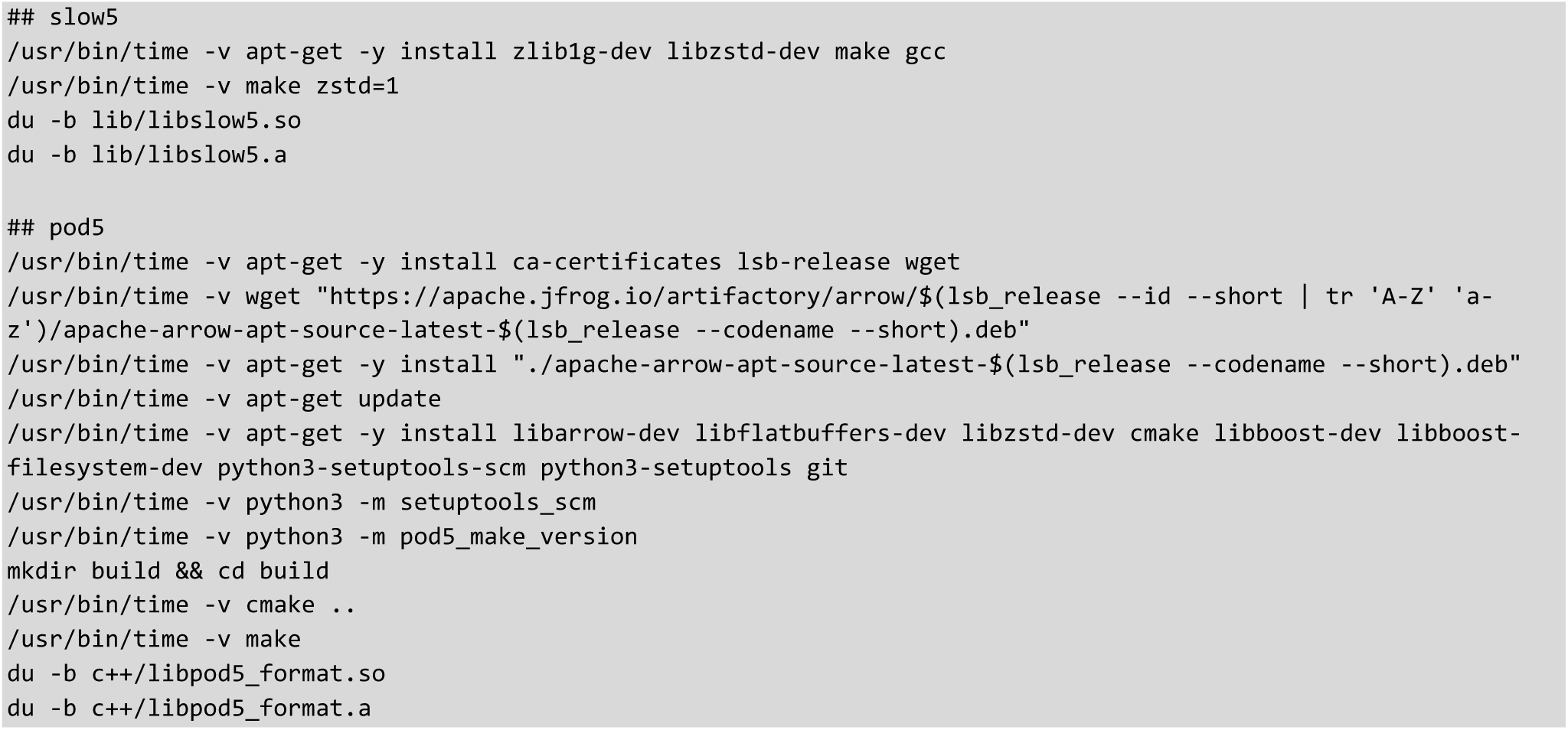

versions:

- slow5lib bench branch [https://github.com/hasindu2008/slow5lib/tree/bench commit c254055ea7811ce3daec541141a8608d5dc15243]
- pod5 v0.3.10
- Docker version 26.1.3; *debian:bookworm-slim* image (Debian 12)

#### Notes

- See inside the *docker_slow5.sh* under the *misc* directory of the *slow5-pod5-bench* repository for detailed steps of slow5 compilation
- See inside the *docker_pod5.sh* under the *misc* directory of the *slow5-pod5-bench* repository for detailed steps of pod5 compilation
- reported times for compilation are the sum of all steps that were measured by the time command above
- pod5 v0.3.10 was used instead of pod5 v0.3.2 (the version used for benchmarks), because we could not get this v0.3.2 compiled
- The reason for using Docker is that pod5 library needs specific compiler versions and many packages that were tedious to install without root access. slow5lib is also built inside a Docker for fair comparison of the building time.

#### Source code complexity

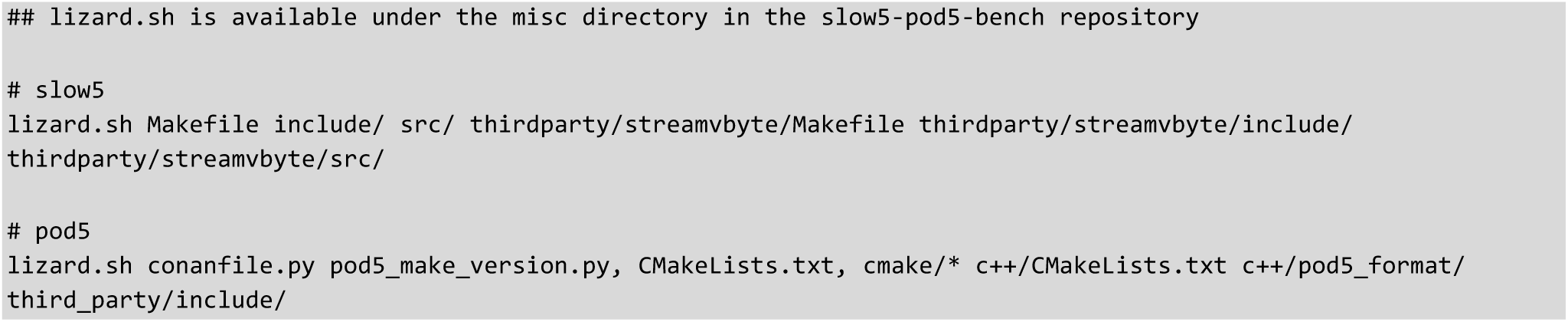

versions:

- slow5: https://github.com/hasindu2008/slow5lib/archive/refs/tags/v1.2.0.tar.gz

pod5: https://github.com/nanoporetech/pod5-file-format/archive/refs/tags/0.3.2.tar.gz

- lizard version 1.17.17 [https://github.com/terryyin/lizard]

Notes:

For slow5, makefile source code is included because that is the preferred option. Optional cmake is not included for slow5. For pod5, no make-based method exist; thus, cmake source code is included. For pod5, we also include conanfile.py and pod5_make_version.py, because they are required to build the C++ version. Source code inside third-party directories is included for both slow5 and pod5, because they contain source inbuilt to the repository, which are not considered external dependencies.

#### Stability/Compatibility

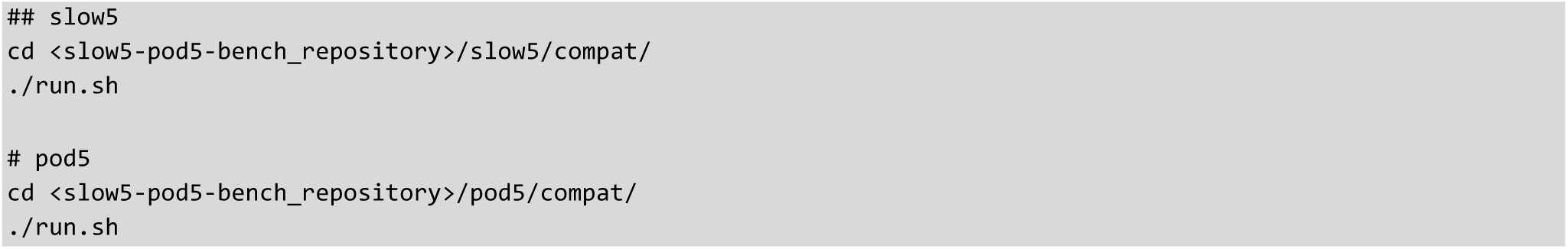

